# Rapid and Sustained Homeostatic Control of Presynaptic Exocytosis at a Central Synapse

**DOI:** 10.1101/785535

**Authors:** Igor Delvendahl, Katarzyna Kita, Martin Müller

## Abstract

Animal behavior is remarkably robust despite constant changes in neural activity. Homeostatic plasticity stabilizes central nervous system (CNS) function on time scales of hours to days. If and how CNS function is stabilized on more rapid time scales remains unknown. Here we discovered that mossy fiber synapses in the mouse cerebellum homeostatically control synaptic efficacy within minutes after pharmacological glutamate receptor impairment. This rapid form of homeostatic plasticity is expressed presynaptically. We show that modulations of readily-releasable vesicle pool size and release probability normalize synaptic strength in a hierarchical fashion upon acute pharmacological and prolonged genetic receptor perturbation. Presynaptic membrane capacitance measurements directly demonstrate regulation of vesicle pool size upon receptor impairment. Moreover, presynaptic voltage-clamp analysis revealed increased calcium-current density under specific experimental conditions. Thus, homeostatic modulation of presynaptic exocytosis through specific mechanisms stabilizes synaptic transmission in a CNS circuit on time scales ranging from minutes to months. Rapid presynaptic homeostatic plasticity may ensure stable neural circuit function in light of rapid activity-dependent plasticity.

## Introduction

Adaptive animal behavior presupposes plastic, and yet stable neural function. While neural activity is constantly changing because of activity-dependent plasticity (1, 2), robust animal behavior can be maintained for a lifetime. However, stable nervous system function and behavior are not a given, as apparent from pathological states, such as epilepsy or migraine (3, 4). A number of studies revealed that homeostatic plasticity actively stabilizes neural excitability and synaptic transmission (5–8). The most widely studied form of homeostatic synaptic plasticity is synaptic scaling – the bidirectional, compensatory regulation of neurotransmitter receptor abundance (6). This postsynaptic type of homeostatic plasticity is typically observed after chronic pharmacological blockade of action potentials (APs) or neurotransmitter receptors in neuronal cell-culture systems (9–11), but also stabilizes AP firing rates after sensory deprivation *in vivo* (12, 13).

There is also evidence for presynaptic forms of homeostatic synaptic plasticity. Presynaptic homeostatic plasticity (PHP) stabilizes synaptic efficacy at neuromuscular synapses after acute or sustained neurotransmitter receptor perturbation in various species (14–18). In the mammalian CNS, PHP is mostly studied in cell-culture systems after prolonged perturbation of neural activity (8). The expression of PHP depends on cell-culture age, with cultures older than two weeks compensating for prolonged activity deprivation through concomitant regulation of presynaptic release, postsynaptic receptor abundance, and synapse number (19–22). Despite evidence for compensatory changes in presynaptic structure and function (23–27), it is currently not well understood how these relate to the postsynaptic modifications during synaptic scaling. Moreover, it has remained largely elusive if PHP stabilizes the activity of native neural circuits (8, 28, 29).

The maintenance of robust CNS function is strongly challenged by Hebbian plasticity, which is induced on rapid time scales (2) and eventually destabilizes neuronal activity (30–32). In contrast, the expression of pre- and postsynaptic homeostatic plasticity requires prolonged perturbation of CNS function for hours to days (8), raising the question if and how CNS function is stabilized on rapid time scales. Given the discrepancy in temporal dynamics, homeostatic plasticity is thought to slowly integrate and normalize neural activity changes that happen on faster time scales (7, 11). However, theoretical work strongly implies that the slow time course of homeostatic plasticity is insufficient to prevent instabilities induced by Hebbian plasticity (33).

Here we define the time course of a fast, presynaptic form of homeostatic plasticity at a cerebellar synapse. Employing direct pre- and postsynaptic whole-cell patch clamp recordings in acute brain slices, we discover that a synergistic modulation of readily-releasable vesicle pool size and vesicular release probability through elevated Ca^2+^-channel density underlies the homeostatic control of exocytosis from these CNS terminals. Rapid PHP may stabilize synaptic information transfer to balance changes imposed by perturbations or Hebbian plasticity.

## Results

### Rapid, Reversible Homeostatic Release Modulation

We probed if fast presynaptic homeostatic mechanisms stabilize synaptic transmission at central synapses in an acute mouse brain slice preparation. To this end, we recorded miniature- (mEPSCs) and AP-evoked excitatory postsynaptic currents (EPSCs) at adult cerebellar mossy fiber (MF)-granule cell (GC) synapses (Fig. 1A) (34, 35) after incubation with the non-competitive α-amino-3-hydroxy-5-methyl-4-isoxazolepropionic acid receptor (AMPAR) blocker GYKI 53655 for various incubation times (see *Methods*). GYKI application at sub-saturating concentration (2 µM, (36)) for 5–20 minutes reduced mEPSC and evoked EPSC amplitudes by ∼30% without affecting mEPSC and EPSC kinetics (Figs. 1B, 1C, *SI Appendix* Figs. S1A–G). By contrast, recordings obtained after 20 minutes of GYKI incubation revealed EPSC amplitudes that were comparable to control levels (Figs. 1B–D). Similar results were obtained after longer incubation times of up to 100 minutes (Figs. 1B–D). Because GYKI application equally reduced mEPSC amplitudes at all incubation times, there was a significant increase in quantal content after 20 minutes of GYKI exposure (Figs. 1C, D), indicative of enhanced neurotransmitter release. The increase in quantal content scaled with the degree of mEPSC reduction, such that EPSC amplitudes were precisely maintained at control levels (*SI Appendix* Fig. S1H), consistent with the phenomenology of PHP at the NMJ (17, 37). These data demonstrate a rapid form of PHP at synapses in the mammalian CNS upon acute, sub-saturating AMPAR blockade on the minute time scale.

**Fig. 1:**
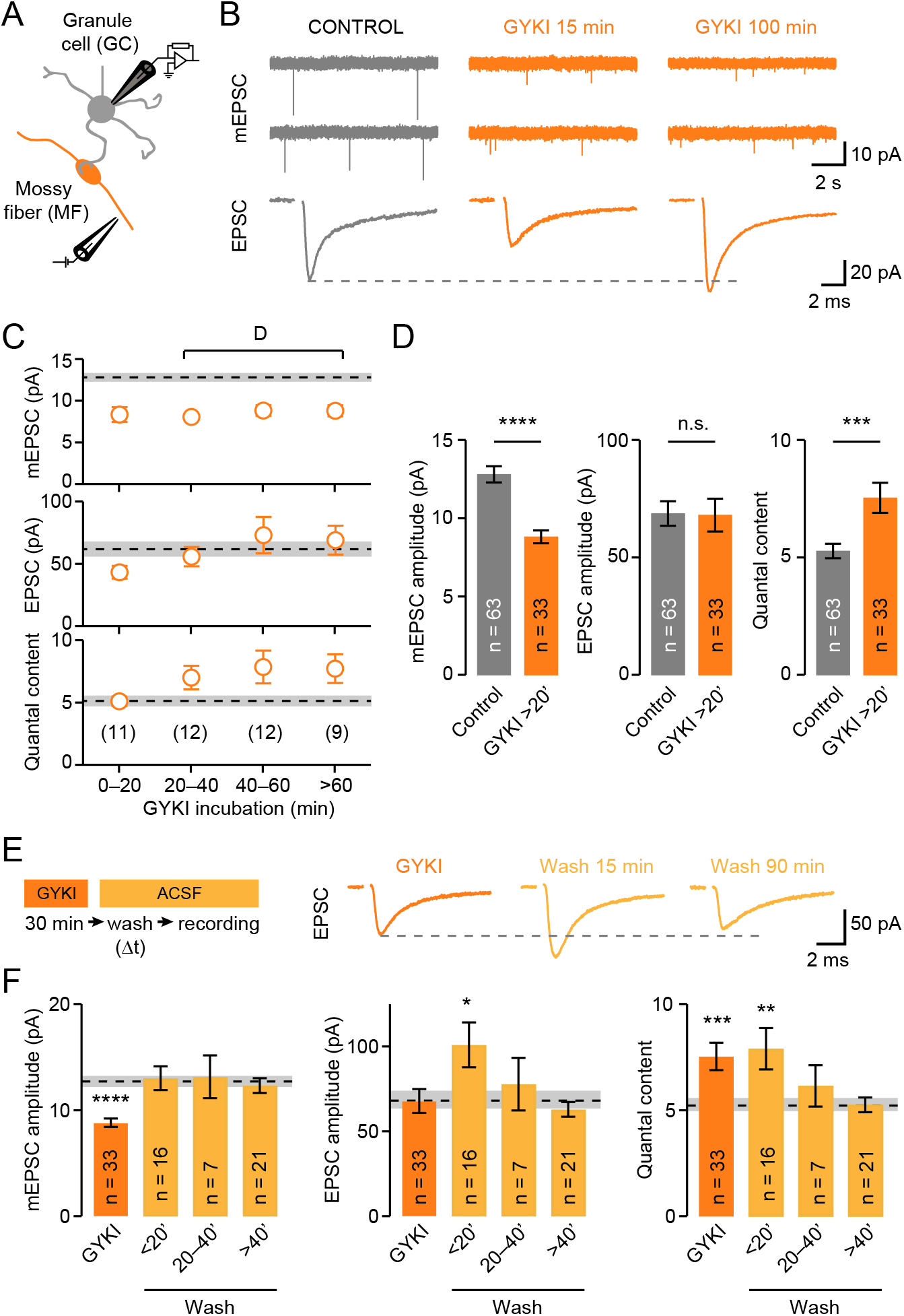
Rapid, Reversible Homeostatic Release Modulation in the Mouse Cerebellum. (A) Schematic of postsynaptic whole-cell recordings from cerebellar granule cells and MF stimulation to evoke EPSCs. (B) Representative mEPSCs and AP-evoked EPSCs recorded from a control synapse (Left), or from synapses incubated with 2 µM of the AMPAR antagonist GYKI 53655 for the indicated incubation times (Middle and Right). (C) mEPSC amplitudes, EPSC amplitudes, and quantal content for different GYKI incubation times. Dashed lines represent control values with shaded area indicating ± SEM, numbers represent individual recordings. (D) Average data for mEPSC amplitude (Cohen’s *d* = −1.1; p = 1.7E−6), EPSC amplitude (*d* = −0.02; p = 0.94) and quantal content (= EPSC/mEPSC; *d* = 0.78; p = 0.0005) of all cells with >20 minutes of GYKI incubation. (E) Left: Experimental design. Slices were incubated in ACSF containing 2 µM GYKI for 30 minutes and recordings were subsequently carried out in control ACSF (“wash”) at indicated time intervals (Δt). Right: example EPSCs for GYKI and wash conditions. (F) Average data for mEPSC amplitude, EPSC amplitude, and quantal content. The group ‘wash <20 minutes’ includes cells with wash-out during recording. Dashed lines indicate control averages with shaded area representing ± SEM. * p < 0.05; ** p < 0.01; *** p < 0.001; **** p < 0.0001; n.s. not significant; two-tailed Student’s t-test. Data are presented as mean ± SEM.

Next, we recorded from MF-GC synapses following wash-out of GYKI to investigate the reversibility of PHP (Fig. 1E; see *Methods*). Slices were incubated with GYKI for 30 minutes and then exposed to control ACSF for varying times before obtaining whole-cell recordings. GYKI wash-out increased mEPSC and EPSC amplitudes, with quantal content remaining significantly larger than under control conditions within ∼20 minutes after wash-out (Fig. 1F). This implies that PHP expression persists when AMPAR perturbation is removed on a short time scale. Additional recordings demonstrated that PHP was fully reversible within ∼20–40 minutes after GYKI wash-out (Fig. 1F), similar to previous reports at the mouse NMJ (18). Thus, PHP completely reverses within tens of minutes after reattained receptor function.

### Acute Homeostatic RRP Size Modulation

To assess the presynaptic mechanisms underlying neurotransmitter release potentiation upon AMPAR perturbation, we first tested if changes in the number of readily-releasable vesicles (readily-releasable pool, RRP) (38) contribute to PHP following GYKI application. To estimate RRP size, we used high-frequency train stimulation and analysis of cumulative EPSC amplitudes (Fig. 2A) (39). This analysis gives an effective RRP size estimate assuming that EPSC amplitudes depress because of vesicle pool depletion. 300-Hz stimulation revealed a pronounced increase in RRP size in the presence of GYKI (Fig. 2B, *SI Appendix* Figs. S3C–E) that paralleled the time course of PHP induction and reversal (*SI Appendix* Figs. S3A–B). An increase in release probability (*p_r_*) could also contribute to release potentiation during PHP. However, *p_r_* estimated from EPSC train stimulation was unchanged after GYKI incubation (Fig. 2B). Moreover, EPSC amplitude coefficient of variation (CV; *SI Appendix* Fig. S3F) and EPSC paired-pulse ratios (PPRs; Fig. 2D; PPRs were recorded under conditions of minimized postsynaptic contributions (40); *SI Appendix* Fig. S3G), which are inversely correlated with *p_r_* (41), were unaffected by GYKI treatment, indicating that *p_r_* was largely unchanged.

**Fig. 2:**
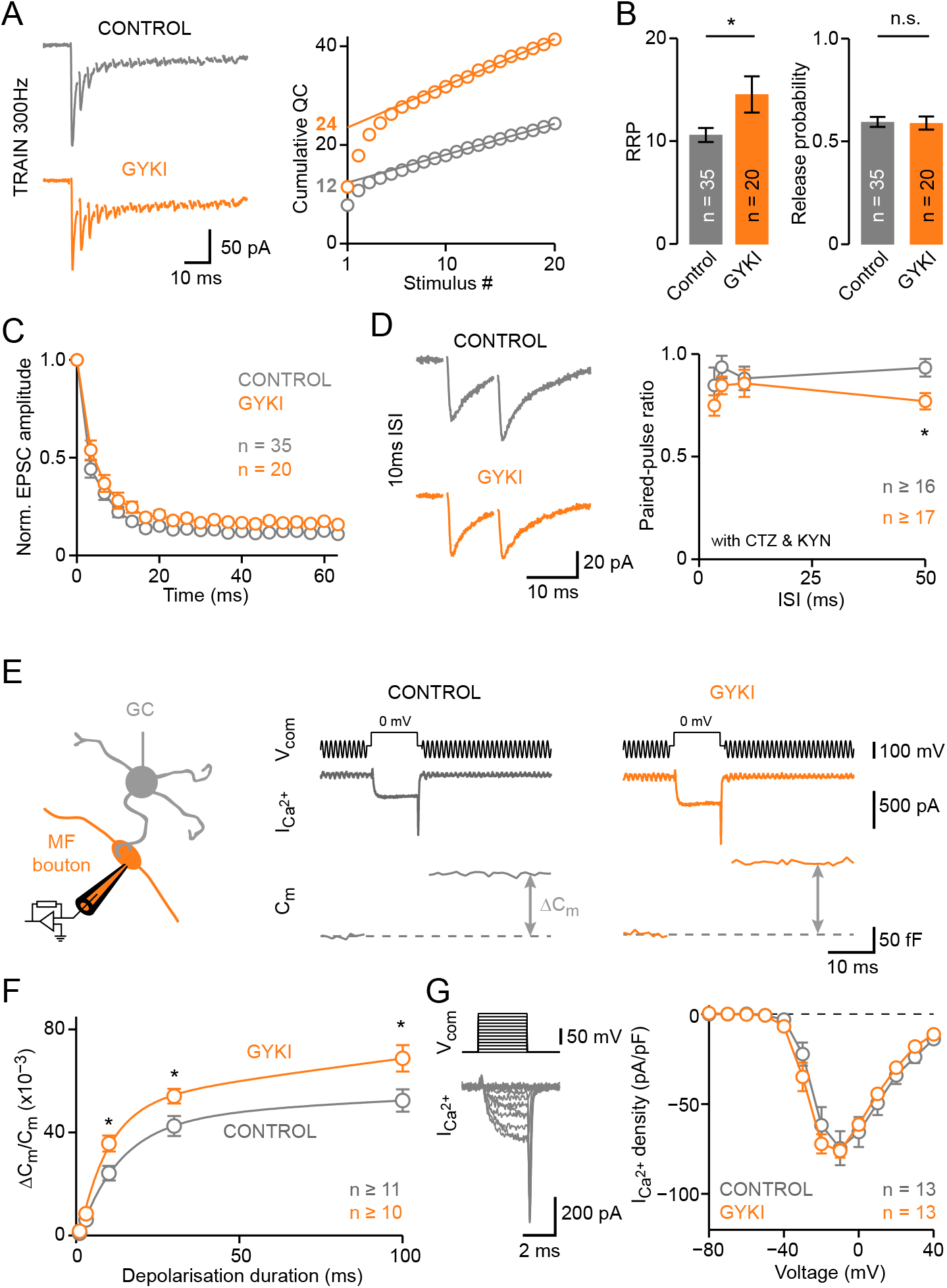
Acute AMPAR Perturbation Increases Readily-Releasable Pool Size. (A) Left: Representative recordings of 300-Hz stimulation (20 pulses) for control synapses and GYKI-treated synapses (>20 minutes incubation). Right: Cumulative quantal content [calculated as: (cumulative EPSC amplitude)/(mEPSC amplitude)] for the examples on the left. Extrapolated linear fits provide the RRP size estimate (indicated). (B) Average data for control and GYKI. GYKI enhanced RRP size (*d* = 0.69; p = 0.017) without changing release probability [calculated as: (first EPSC amplitude)/(cumulative EPSC amplitude); *d* = −0.04; p = 0.89]. (C) Average EPSC amplitude during 300-Hz train stimulation for control and GYKI, normalized to the first EPSC. (D) Left: Representative EPSCs evoked by stimulation with 10 ms inter-stimulus interval (ISI) in the presence of cyclothiazide and kynurenic acid to minimize postsynaptic contributions. Right: Average paired-pulse ratio (PPR) versus ISI for control and GYKI-treated cells. PPRs were not affected by GYKI treatment. (E) Left: Cartoon illustrating presynaptic whole-cell recordings from cerebellar mossy fiber boutons. Right: Voltage command (V_com_, 10-ms depolarization to 0 mV, top), representative pharmacologically isolated Ca^2+^ currents (middle), and membrane capacitance (C_m_) jumps (bottom) for control and GYKI-treated boutons (>20 minutes incubation). (F) Average data of C_m_ increase (ΔC_m_) versus duration of presynaptic depolarization for both conditions (η^2^ = 0.14; p = 6.6E−05; two-way ANOVA). Lines are bi-exponential fits; C_m_ increase was normalized to resting whole-cell capacitance. (G) Left: Representative recording of pharmacologically isolated presynaptic Ca^2+^ currents. Ca^2+^ currents were evoked by 3-ms voltage steps from a holding potential of −80 mV to +40 mV in 10-mV increments. Right: Ca^2+^-current density versus voltage for control and GYKI (η^2^ = <0.01; p = 0.88; two-way ANOVA). * p < 0.05; ** p < 0.01; *** p < 0.001; **** p < 0.0001; n.s. not significant; two-tailed Student’s t-test. Data are presented as mean ± SEM.

To directly probe presynaptic function during pharmacological AMPAR perturbation, we performed presynaptic whole-cell recordings from MF boutons (Fig. 2E) (35, 42). First, we employed membrane capacitance (C_m_) measurements (43, 44) to study exocytosis. Brief depolarizations under voltage-clamp caused Ca^2+^ influx and a jump in C_m_ corresponding to synaptic vesicle exocytosis (Fig. 2E) (35, 45–47) and providing a direct estimate of RRP size. MF boutons displayed significantly larger C_m_ jumps upon depolarizations of various durations in the presence of GYKI (>20 minutes incubation) compared to untreated controls (Fig. 2F). The enhanced exocytosis upon GYKI treatment corroborates the increase in effective RRP size estimated from postsynaptic recordings (Fig. 2B). Our presynaptic capacitance data provide an independent confirmation of changes in RRP size, because postsynaptic estimates of RRP are based on relating evoked EPSCs to mEPSCs, which may involve partially distinct vesicle pools and/or postsynaptic receptors (48, 49) (cf. *SI Appendix* Fig. S2). Whereas absolute RRP size estimates obtained from presynaptic C_m_ measurements and postsynaptic EPSC analysis differ because of the high bouton-to-granule cell connectivity (35, 50), we observed a similar relative RRP increase after GYKI treatment in pre- and postsynaptic recordings (∼35%; Figs. 2B and 2F).

Next, we investigated pharmacologically-isolated presynaptic whole-cell Ca^2+^-currents after GYKI application (Fig. 2G). Presynaptic Ca^2+^ currents were elicited by 3-ms voltage steps, and steady-state amplitudes, as well as tail currents were analyzed. Ca^2+^-current density (Fig. 2G), steady-state activation, and activation or deactivation time constants were similar between GYKI-treated and untreated MF boutons (*SI Appendix* Fig. S3I), indicating no major differences in Ca^2+^-channel levels, voltage dependence of Ca^2+^-channel activation, or channel kinetics. No apparent changes in presynaptic Ca^2+^ influx at GYKI-treated terminals indicate largely unaltered *p_r_*, in line with train-based *p_r_* estimates (Fig. 2B–C) and PPR data (Fig. 2D). Together, these data establish that acute AMPAR perturbation at a mammalian central synapse leads to enhanced presynaptic exocytosis and a rapid and reversible expansion of RRP size without apparent *p_r_* changes.

### Sustained Homeostatic RRP Size Modulation

To investigate if PHP can be sustained over longer time periods, we recorded from adult heterozygous mice carrying a loss-of-function mutation in *GRIA4*, the gene encoding the GluA4 AMPAR-subunit (51, 52). GluA4 is expressed in cerebellar GCs (36, 53) and confers a large conductance to synaptic AMPARs (54). Importantly, heterozygous *GRIA4* mice, henceforth called “*GluA4^+/−^*“, have reduced GluA4 protein levels (52, 55). *GluA4^+/−^* synapses had significantly smaller mEPSC amplitudes (Fig. 3A–B, *SI Appendix* Figs. S4A–B and S5). In contrast, EPSC amplitudes were similar between *GluA4^+/−^* synapses and controls (Fig. 3A–B), translating into a significant increase in quantal content (Fig. 3B). We next probed RRP size at *GluA4^+/−^* synapses. 300-Hz train stimulation revealed a significant increase in effective RRP size at *GluA4^+/−^* synapses (Fig. 3C–D, *SI Appendix* Fig. S4F), similar to the results seen after GYKI application to WT synapses (Fig. 2A–B). We did not observe significant changes in *p_r_* estimated from stimulus trains (Fig. 3D), PPR (*SI Appendix* Fig. S4D), or EPSC amplitude CV (*SI Appendix* Fig. S4E) between *GluA4^+/−^* and WT synapses. Taken together, these results provide evidence for increased neurotransmitter release and RRP size after chronic AMPAR impairment.

**Fig. 3:**
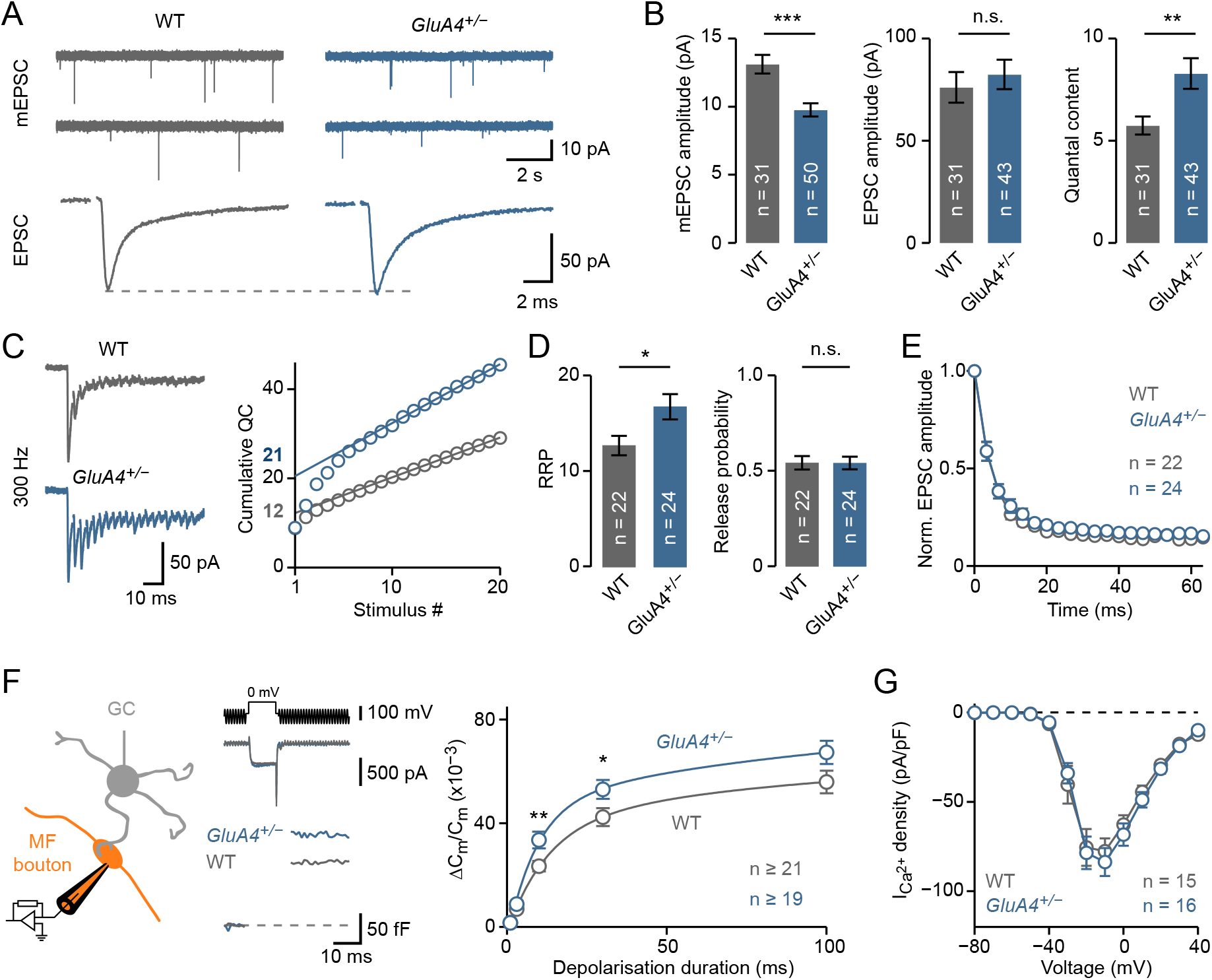
Sustained AMPAR Perturbation Increases Readily-Releasable Pool Size. (A) Representative mEPSCs and AP-evoked EPSCs in WT and *GluA4^+/−^* mice. (B) Average data for mEPSC amplitude (*d* = −0.90; p = 0.0001), EPSC amplitude (*d* = 0.14; p = 0.55) and quantal content (= EPSC/mEPSC; *d* = 0.70; p = 0.0095). (C) Left: Representative recordings after 300-Hz stimulation (20 pulses) for WT and *GluA4^+/−^*. Right: Cumulative quantal content [calculated as: (cumulative EPSC amplitudes)/(mEPSC amplitude)] for the examples on the left. Extrapolated linear fits provide the RRP size estimate (indicated). (D) Average data for WT and *GluA4^+/−^*. RRP size was increased (*d* = 0.72; p = 0.02) at *GluA4^+/−^* synapses without apparent changes in release probability [calculated as: (first EPSC amplitude)/(cumulative EPSC amplitude); *d* = −0.001; p = 0.99]. (E) Average normalized EPSC amplitude during 300-Hz train stimulation. (F) Left: Schematic of recordings from MF boutons. Middle: Voltage command (V_com_, 10-ms depolarization to 0 mV, top), representative Ca^2+^ currents (middle) and membrane capacitance (C_m_) jumps (bottom) for WT and *GluA4^+/−^*. Right: Average presynaptic C_m_ increase (ΔC_m_) against stimulus duration in WT and *GluA4^+/−^* MF boutons (η^2^ = 0.07; p = 0.0002; two-way ANOVA). (G) Ca^2+^-current density versus voltage in WT and *GluA4^+/−^* (η^2^ = 0.001; p = 0.60; two-way ANOVA). * p < 0.05; ** p < 0.01; *** p < 0.001; n.s. not significant; two-tailed Student’s t-test. Data are presented as mean ± SEM.

We then employed presynaptic recordings to study exocytosis and Ca^2+^ influx after genetic AMPAR perturbation. Presynaptic C_m_ measurements revealed enhanced exocytosis in *GluA4^+/−^* MF boutons compared to WT (Fig. 3F), in agreement with the observed increase in effective RRP size based on postsynaptic recordings (cf. Fig. 3D). Furthermore, Ca^2+^-current density was similar in both genotypes (Fig. 3G), consistent with the *p_r_* estimates from postsynaptic recordings. Collectively, these results show that a sustained increase in RRP size—but not *p_r_*— underlies PHP expression upon genetic AMPAR perturbation at MF-GC synapses. As we recorded from adult synapses, our data establish that PHP can be chronically expressed in the mammalian CNS for months.

### Synergistic Homeostatic Control of RRP Size and Release Probability

Next, we investigated PHP after combined pharmacological and genetic AMPAR perturbation. Application of 2 µM GYKI 53655 to slices from *GluA4^+/−^* mice further reduced mEPSC amplitudes by 22%, but EPSC amplitudes remained similar to baseline values (Fig. 4A–B, *SI Appendix* Fig. S5A–E). Accordingly, there was an additional 26% increase in quantal content at *GluA4^+/−^* synapses incubated with GYKI (Fig. 4B). High-frequency train stimulation indicated no additional increase in effective RRP size at GYKI-treated *GluA4^+/−^* synapses (Fig. 4C, *SI Appendix* Fig. S5F), suggesting that another process enhanced release. Indeed, the stimulus train-based *p_r_* estimate was significantly increased upon GYKI treatment (Fig. 4C). Furthermore, there was a significant decrease in EPSC amplitude CV and PPR at *GluA4^+/−^* synapses incubated with GYKI (Fig. 4E–F, *SI Appendix* Fig. S5G), indicating increased *p_r_*. We also observed a correlation between *p_r_* and mEPSC amplitude reduction at GYKI-treated *GluA4^+/−^* synapses, suggesting that *p_r_* potentiation scales with receptor impairment (*SI Appendix* Fig. S6).

**Fig. 4:**
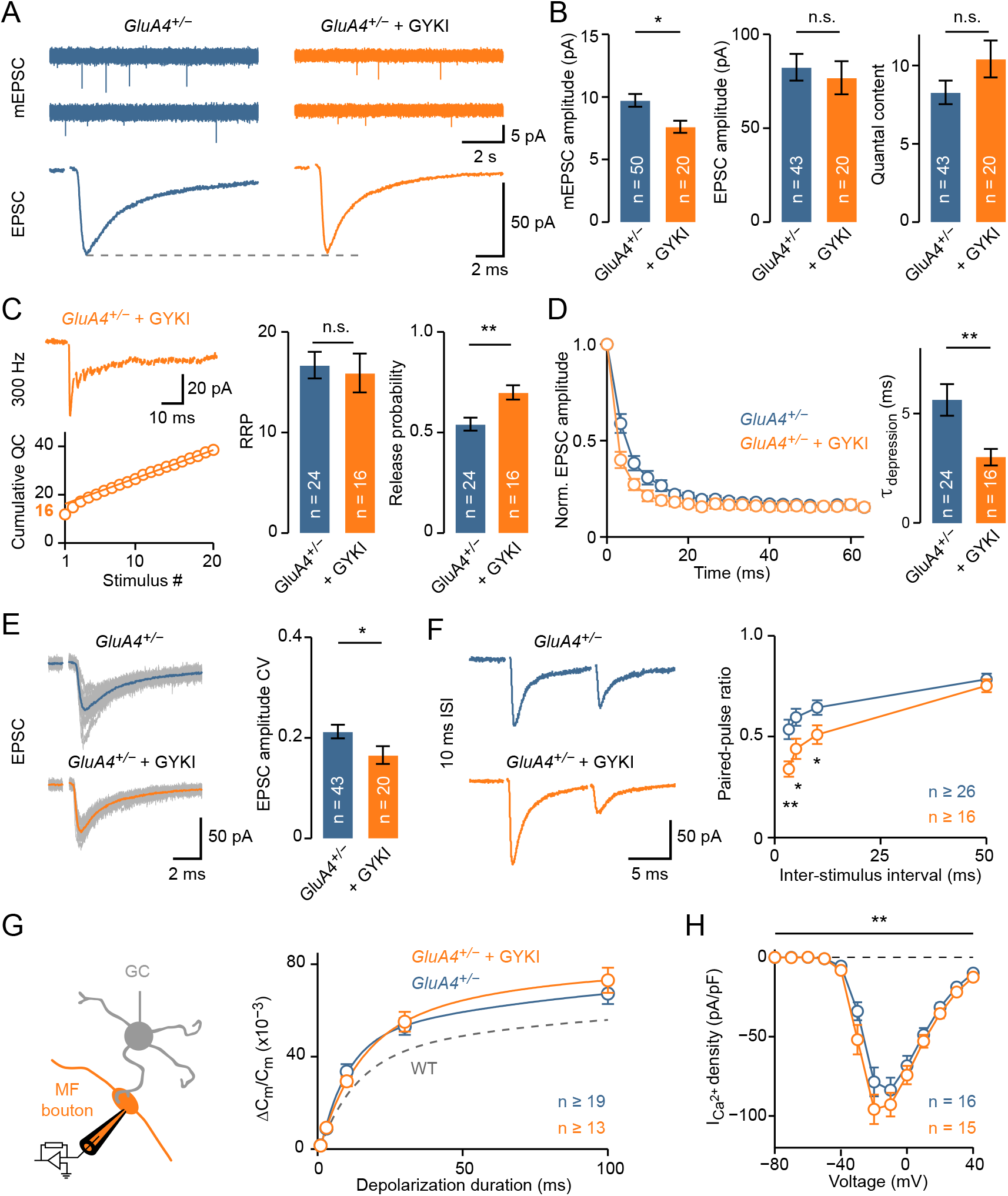
Synergistic Homeostatic Control of RRP Size and Release Probability. (A) Representative mEPSCs and AP-evoked EPSCs in *GluA4^+/−^* mice under control and in the presence of 2 µM GYKI 53655. (B) Average data for mEPSC amplitude (*d* = −0.84; p = 0.011), EPSC amplitude (*d* = −0.14; p = 0.65) and quantal content (= EPSC/mEPSC; *d* = 0.41; p = 0.12) in *GluA4^+/−^* with or without GYKI. (C) High-frequency train-based RRP estimation. Left: Representative 300-Hz train recording in *GluA4^+/−^* with GYKI. Right: Average data for RRP size (*d* = −0.11; p = 0.73) and release probability [calculated as: (first EPSC amplitude)/(cumulative EPSC amplitude); *d* = 1.04; p = 0.003]. (D) Left: Average EPSC amplitude during 300-Hz train stimulation for *GluA4^+/−^* and *GluA4^+/−^* with GYKI, normalized to the first EPSC. Right: Average time constant of synaptic depression during 300-Hz train stimulation (*d* = −0.90; p = 0.0081). Faster depression of GYKI-treated *GluA4^+/−^* synapses suggests increased *p_r_*. (E) Left: Example EPSCs (24 single responses are overlaid with the average) in *GluA4^+/−^* under control and with GYKI. Same examples as in A. Right: Average data of EPSC amplitude coefficient of variation (CV), which was decreased by GYKI (*d* = −0.58; p = 0.04). (F) Left: Representative recordings at 10 ms ISI for *GluA4^+/−^* and *GluA4^+/−^* with GYKI. Right: Average paired-pulse ratios for *GluA4^+/−^* and *GluA4^+/−^* with GYKI. (G) Left: Schematic of presynaptic recordings. Right: Average capacitance jumps normalized to resting C_m_. Direct presynaptic recordings reveal no further enhancement of RRP in *GluA4^+/−^* upon GYKI treatment (η^2^ < 0.001; p = 0.92; two-way ANOVA). Dashed line replots WT data from Fig. 3F for comparison. (H) GYKI slightly enhanced Ca^2+^-current density in *GluA4^+/−^* (η^2^ = 0.02; p = 0.004; two-way ANOVA; post-hoc n.s.). * p < 0.05; ** p < 0.01; *** p < 0.001; n.s. not significant; two-tailed Student’s t-test. Data are presented as mean ± SEM.

Presynaptic capacitance measurements revealed similar C_m_ jumps between GYKI-treated and untreated *GluA4^+/−^* MF boutons (Fig. 4G), indicating no further increase in RRP size with respect to WT (Fig. 4G). Interestingly, there was a small, but significant increase in Ca^2+^-current density in *GluA4^+/−^* MF boutons treated with GYKI (Fig. 4H, cf. Fig. 6D) without apparent changes in Ca^2+^-current kinetics (*SI Appendix* Fig. S5H–I). This suggests elevated presynaptic Ca^2+^-channel levels at GYKI-treated *GluA4^+/−^* terminals. Given the power relationship between presynaptic Ca^2+^ influx and exocytosis (56), the relatively moderate increase in Ca^2+^-current density is expected to produce a significant increase in *p_r_*, in agreement with the enhanced *p_r_* inferred from postsynaptic recordings (Fig. 4C–F). Together, these findings demonstrate that RRP size and *p_r_* can be synergistically modulated to achieve homeostatic stabilization of synaptic transmission, depending on the magnitude or type of receptor perturbation. By extension, this ‘sequential’ regulation of RRP size and *p_r_* points towards a potential hierarchy in the modulation of presynaptic function during PHP (see *Discussion*).

### GluA4 is the Major AMPAR Subunit at MF-GC Synapses

Loss of one *GluA4* copy was sufficient to reduce mEPSC amplitude, indicating an important role for this AMPAR subunit in synaptic transmission at MF-GC synapses. To further elucidate the contribution of GluA4, we recorded from homozygous *GluA4^−/−^* mice that completely lack the large conductance GluA4 AMPAR-subunit (51). There were almost no detectable spontaneous mEPSCs at *GluA4^−/−^* GCs (Fig. 5A), preventing an amplitude quantification. Evoked EPSCs were very small (18% of WT), but could reliably be observed (Fig. 5A–B). The EPSC decay time constant was slower than in WT (Fig. 5C), owing to the loss of the fast GluA4 subunit (52, 57). Thus, GluA4 is the major AMPAR subunit mediating the fast excitation of cerebellar GCs, the most abundant neuronal cell type in the mammalian CNS.

**Fig. 5:**
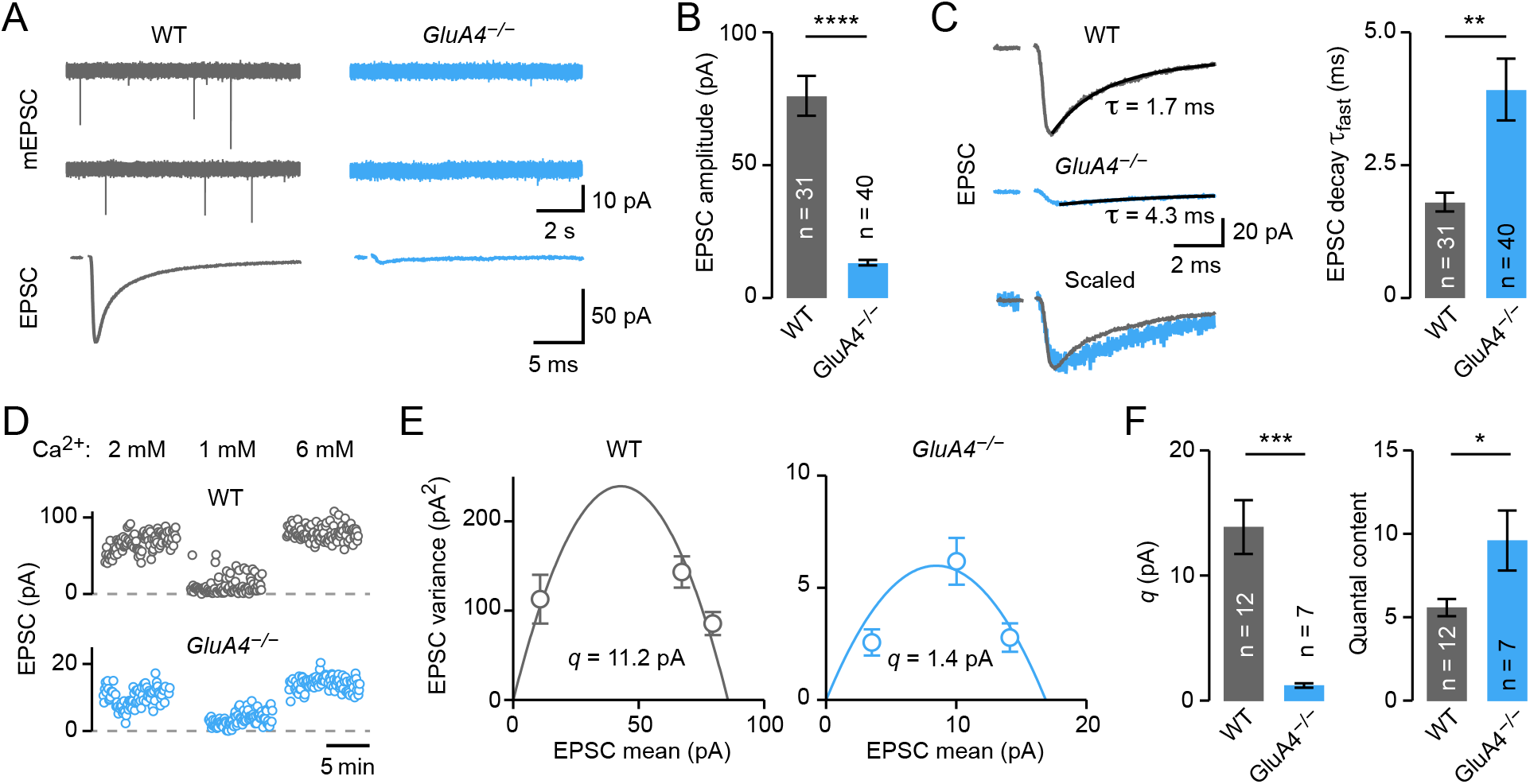
GluA4 is the Major AMPAR Subunit at MF-GC Synapses. (A) Representative mEPSCs and AP-evoked EPSCs in WT and *GluA4^−/−^* mice. (B) Average EPSC amplitudes (*d* = −1.96; p = 8.1E−14). (C) Left: Representative EPSCs in WT and *GluA4^−/−^* mice overlaid with bi-exponential fits to the decay. Time constants of the fast component are indicated. Right: Average fast decay time constant for both genotypes (*d* = 0.84; p = 0.0026). (D) EPSC amplitudes at indicated extracellular Ca^2+^ concentrations of a representative recording in WT (Top) and *GluA4^−/−^* (Bottom). (E) EPSC amplitude variance versus mean for the examples in panel D. Quantal size (*q*) estimates obtained from parabolic fits are indicated. (F) Average data from variance-mean analysis. *GluA4^−/−^* synapses have a strongly reduced *q* (*d* = −2.84; p = 0.0004). Although EPSC amplitudes are smaller in *GluA4^−/−^* (*d* = −2.1; p = 0.0051), quantal content is increased (*d* = 1.01; p = 0.017). * p < 0.05; ** p < 0.01; *** p < 0.001; two-tailed Student’s t-test. Data are presented as mean ± SEM.

The pronounced reduction in EPSC amplitude at MF-GC synapses lacking two *GluA4* copies suggests that PHP is not engaged, saturated, or masked by too strong AMPAR impairment. To distinguish between these possibilities, we first asked if GluA4-lacking synapses express PHP. The small amplitude of mEPSCs (Fig. 5A) made quantification impossible and hence precluded direct calculation of quantal content. We therefore used multiple-probability fluctuation analysis (40, 58) at MF-GC synapses from WT and *GluA4^−/−^* mice to dissect quantal parameters of synaptic transmission. We recorded EPSCs under several *p_r_* conditions by varying the extracellular Ca^2+^ concentration ([Ca^2+^]_e_) (Fig. 5D) and plotted EPSC amplitude variance against mean EPSC amplitude (Fig. 5E). A parabolic fit to the data provided an estimate for quantal size (*q*), which was 1.2 ± 0.2 pA in *GluA4^−/−^* (9% of WT; Fig. 5F, *SI Appendix* Fig. S7A). Using this *q* estimate and the EPSC amplitude at 2 mM [Ca^2+^]_e_, quantal content was strongly increased in *GluA4^−/−^* mutants compared to WT synapses (172% of WT; Fig. 5F, *SI Appendix* Fig. S7B–E). Thus, even under strong genetic AMPAR perturbation by ablation of two *GluA4* copies, MF-GC synapses show a prominent increase in neurotransmitter release. Yet, this increase in release is not sufficient to maintain EPSC amplitudes at WT levels, suggesting that PHP is saturated or masked by too strong receptor impairment.

### Enhanced Exocytosis and Ca^2+^ Influx at *GluA4^−/−^* Boutons

So far, the evidence for presynaptic modulation at *GluA4^−/−^* synapses is based on postsynaptic recordings that are likely limited by the strong reduction in AMPAR currents. We therefore used presynaptic recordings (Fig. 6A) to directly investigate how neurotransmitter release is modulated at *GluA4^−/−^* boutons, and to ask if PHP is saturated or masked. *GluA4^−/−^* boutons displayed significantly larger C_m_ jumps than WT upon presynaptic depolarizations of various durations (Fig. 6B). The enhanced exocytosis in *GluA4^−/−^* is consistent with the increase in quantal content deduced from postsynaptic recordings (Fig. 5F, *SI Appendix* Fig. S7A). Moreover, C_m_ jumps were larger in *GluA4^−/−^* animals compared to *GluA4^+/−^* (*SI Appendix* Figs. S8C, S8E), suggesting a pronounced increase in RRP size that correlates with the degree of *GluA4* loss. Nevertheless, the increase in ΔC_m_ in *GluA4^−/−^* (∼40%) is at least an order of magnitude smaller than what would be required to maintain WT EPSC levels (cf. Fig. 6B, *SI Appendix* Fig. S7A). We thus conclude that PHP is saturated under these conditions.

**Fig. 6:**
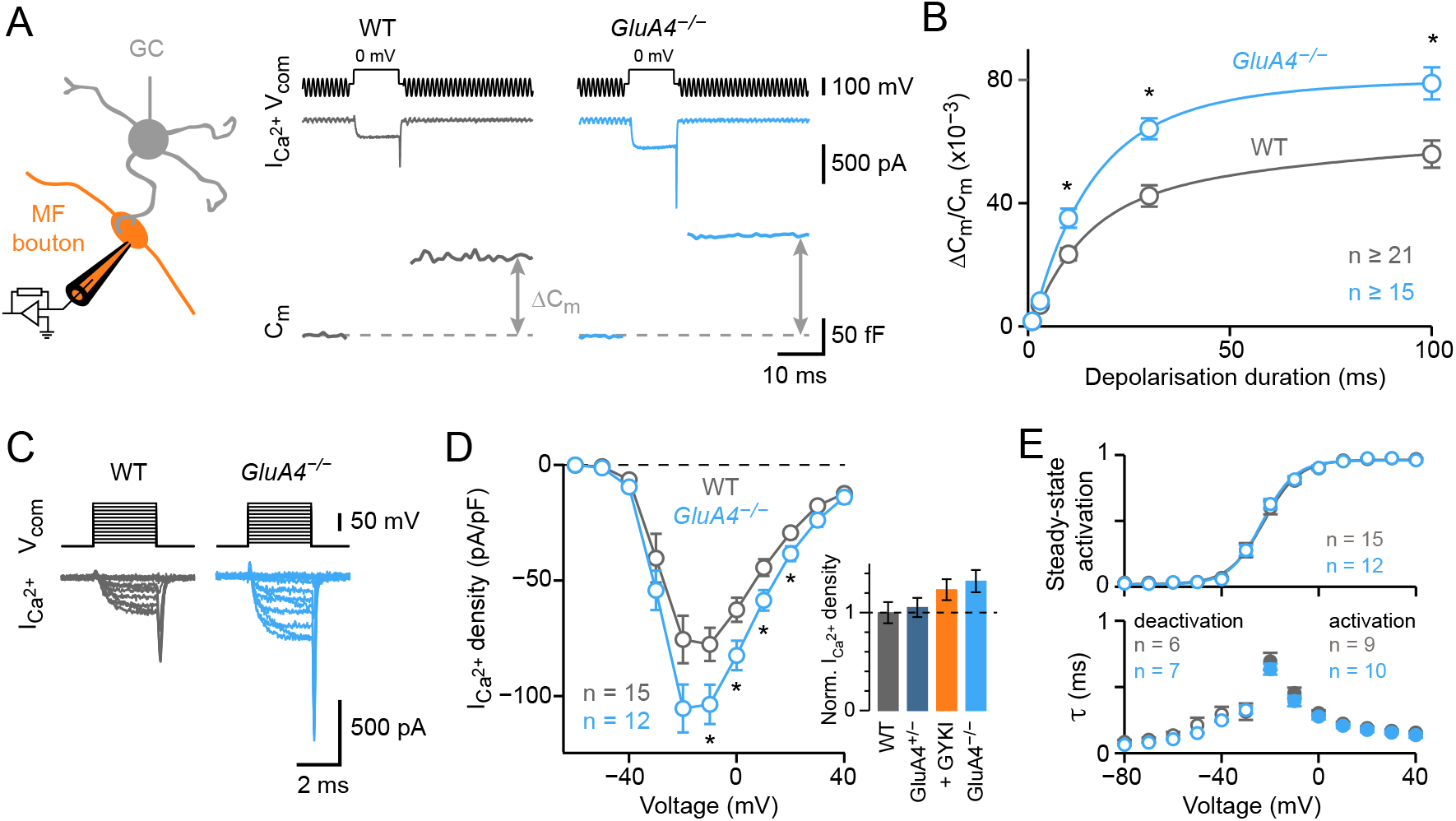
Enhanced Exocytosis and Ca^2+^ Influx at GluA4-Deficient Boutons. (A) Left: Schematic of recordings from MF boutons. Right: Voltage command (V_com_, 10-ms depolarization to 0 mV, top), representative Ca^2+^ currents (middle) and membrane capacitance (C_m_) jumps (bottom) for WT and *GluA4^−/−^*. (B) Average C_m_ increase (ΔC_m_) versus duration of presynaptic depolarization (η^2^ = 0.179; p = 2.2E−09; two-way ANOVA). Lines are bi-exponential fits; C_m_ increase was normalized to resting whole-cell capacitance. (C) Representative pharmacologically isolated presynaptic Ca^2+^ currents. Ca^2+^ currents were evoked by 3-ms voltage steps from a holding potential of −80 mV to +40 mV in 10-mV increments. (D) Ca^2+^-current density versus voltage for WT and *GluA4^−/−^* (η^2^ = 0.065; p = 3.1E−06; two-way ANOVA). Inset shows maximum Ca^2+^-current density of WT, *GluA4^+/−^*, *GluA4^+/−^* with GYKI, and *GluA4^−/−^* for comparison (normalized to WT). (E) Top: Average data for steady-state activation; lines are sigmoidal fits. Bottom: Time constants of Ca^2+^-channel activation (filled symbols) and deactivation (open symbols). * p < 0.05; ** p < 0.01; two-tailed Student’s t-test. Data are presented as mean ± SEM.

To further explore the mechanisms underlying the increase in exocytosis observed at *GluA4^−/−^* synapses, we recorded presynaptic Ca^2+^ currents at MF boutons (Fig. 6C). Ca^2+^-current density of *GluA4^−/−^* boutons was significantly enhanced (Fig. 6D, *SI Appendix* Fig. S8D). There was no change in Ca^2+^-current steady-state activation or time constants of activation and deactivation in *GluA4^−/−^* mutants (Fig. 6E), indicating no major differences in the voltage dependence of Ca^2+^-channel activation or channel kinetics. Presynaptic current-clamp analysis revealed no major differences in resting membrane potential, AP amplitude and AP duration between *GluA4^−/−^* and WT MF boutons (*SI Appendix* Fig. S8B). Collectively, these data suggest that increased presynaptic Ca^2+^ influx due to elevated Ca^2+^-channel density enhances *p_r_* in *GluA4^−/−^* mutants. In combination with the increase in ΔC_m_ (Fig. 6B), this indicates a synergistic modulation of RRP size and *p_r_* in the *GluA4^−/−^* mutant background.

## Discussion

Our results establish that distinct presynaptic homeostatic mechanisms compensate for neurotransmitter receptor impairment on time scales ranging from minutes to months at a mammalian central synapse. In the CNS, pre- and postsynaptic forms of homeostatic plasticity are typically observed after prolonged perturbation of neuronal activity for hours to days (9–11, 23, 24, 59, 60). Thus, the homeostatic control of neurotransmitter release uncovered here is expressed on a comparably fast time scale, similar to neuromuscular synapses in *Drosophila* and mouse (14, 18). We also uncovered that acutely induced PHP fully reverses within tens of minutes. In *Drosophila*, there is recent evidence that PHP is reversible within days, although the reversibility time course was likely limited by the nature of receptor perturbation (61). By contrast, PHP reverses within minutes after the removal of pharmacological receptor inhibition at the mouse NMJ (18), similar to our results. The rapid time course of PHP may allow stabilizing information transfer in the CNS on fast time scales. It is an intriguing possibility that PHP serves as a mechanism to compensate for Hebbian plasticity at CNS synapses. In this regard, our work may have implications for neural network simulations incorporating Hebbian plasticity rules, because their stability requires fast compensatory processes with similar temporal dynamics as the form of PHP discovered here (33).

The major presynaptic mechanism that was modulated upon AMPAR perturbation in every experimental condition is RRP size. In addition to indirect evidence for RRP size regulation based on postsynaptic recordings, we directly demonstrate increased RRP size employing presynaptic C_m_ measurements for all experimental conditions. Importantly, presynaptic RRP measurements are unaffected by potential differences between evoked and spontaneous release (48, 49). Homeostatic regulation of RRP or recycling pool size has been observed at several synapses in different species (18, 20, 62–64), implying a general, evolutionarily-conserved mechanism.

In addition to RRP size, we provide evidence for homeostatic *p_r_* modulation under specific experimental conditions. Homeostatic *p_r_* changes have been described for the *Drosophila* NMJ (65, 66) and in mammalian CNS cell culture (20, 27), suggesting evolutionary conservation. A major factor that has been linked to homeostatic *p_r_* control is the modulation of presynaptic Ca^2+^ influx (24, 65). Consistently, we uncovered elevated Ca^2+^-current density in the experimental conditions with increased *p_r_*, implying increased levels of presynaptic voltage-gated Ca^2+^ channels. This observation is in line with previous light-microscopy based results from mammalian CNS synapses and the *Drosophila* NMJ (23, 66–68).

We only observed presynaptic Ca^2+^ influx and *p_r_* changes in homozygous *GluA4^−/−^* mutants and after pharmacological receptor perturbation in the *GluA4^+/−^* background. The specific modulation under these experimental conditions may be due to the nature or degree of receptor perturbation. The fact that pharmacological inhibition of WT receptors by a similar degree as in *GluA4^+/−^* mutants induces no apparent changes in *p_r_* argues against the latter possibility (*SI Appendix* Fig. S6). Future work is required to explore the links between receptor perturbation and *p_r_* modulation during PHP.

Whereas RRP size changed in every experimental condition, *p_r_* modulation was only seen in combination with RRP regulation, suggesting a synergistic, hierarchical modulation of RRP size and *p_r_* during PHP. There is no coherent picture regarding the relationship between RRP and *p_r_* regulation during PHP at other synapses. At the *Drosophila* NMJ, genetic data suggests that distinct molecular pathways underlie homeostatic modulation of RRP and Ca^2+^ influx/*p_r_* (64). There is recent evidence that homeostatic vesicle pool regulation requires altered Ca^2+^ influx through P/Q-type Ca^2+^ channels at cultured mammalian synapses (24). By contrast, PHP at the mouse NMJ involves RRP modulation without any evidence for changes in *p_r_* (18). Although future experiments are required to relate these observations, this points towards the possibility that different synapses may engage different presynaptic mechanisms during PHP.

A major finding of this study is that synaptic transmission at cerebellar MF-GC strongly depends on the GluA4 AMPAR subunit. The GluA4 subunit, encoded by *GRIA4*, confers rapid kinetics and large conductance to mammalian AMPARs (53). Previous work at the calyx of Held synapse demonstrated a role of GluA4 for fast sensory processing and indicated partial presynaptic compensation in *GluA4^−/−^* mice (57). Cerebellar mossy fibers are capable of firing at remarkably high frequencies (35) and our data show that GluA4 is the predominant AMPAR subunit at MF-GC synapses. These findings suggest that different CNS synapses rely on the GluA4 subunit for rapid information transfer. Mutations in *GRIA4* have been implicated in schizophrenia and intellectual disability (69, 70), underscoring the importance of this AMPAR subunit for synaptic function.

Our findings advance our understanding of homeostatic plasticity by revealing rapid and reversible regulation of specific presynaptic mechanisms upon glutamate receptor perturbation at a central synapse. In addition, we demonstrate a conserved process between presynaptic homeostatic plasticity studied at the *Drosophila* NMJ and the maintenance of neural function in the mammalian CNS. It will be exciting to explore the underlying molecular mechanisms and the interface between PHP and other forms of synaptic plasticity in the CNS (32, 33, 71).

## Methods

### Animals

Animals were treated in accordance with national and institutional guidelines. All experiments were approved by the Cantonal Veterinary Office of Zurich (authorization no. ZH206/16). Wild-type C57BL/6JRj mice were obtained from Janvier Labs; *GRIA4* knock-out mice were a kind gift of H. Monyer ((51); termed “*GluA4^+/−^*“ and “*GluA4^−/−^*“ throughout the manuscript). *GluA4^−/−^*, *GluA4^+/−^*, and wild-type (“WT”) littermates were bred from heterozygous crosses. Adult (3–10-week-old) mice of either sex were used for experiments. All animals were maintained with food ad libitum on a 12h/12h light/dark cycle, and experiments were performed between 9AM and 9PM.

Genotyping of *GRIA4* mice was performed by PCR analysis of genomic DNA from toe biopsies. For genotyping, the forward primer 5’-CGTGCGCCACCACCGCCCGG-3’ was used with the reverse primer 5’-TGCCACTCAGTTATTGCATCAC-3’ to detect a 290 bp band in wild-type, and with the reverse primer 5’-CAAAACATGGATTAGTCTTTATGGAACAG-3’ to detect a 500 bp band in *GRIA4* mutant mice.

### Slice Electrophysiology

Mice were sacrificed by rapid decapitation; the cerebellar vermis was removed quickly and mounted in a chamber filled with cooled extracellular solution. 300-µm thin parasagittal slices were cut using a Leica VT1200S vibratome (Leica Microsystems), transferred to an incubation chamber at ∼35 °C for 30 minutes and then stored at room temperature until experiments. The extracellular solution (artificial cerebrospinal fluid, ACSF) for slice cutting and storage contained (in mM): 125 NaCl, 25 NaHCO_3_, 20 glucose, 2.5 KCl, 2 CaCl_2_, 1.25 NaH_2_PO_4_, 1 MgCl_2_, equilibrated with 95% O_2_ and 5% CO_2_, pH 7.3, ∼310 mOsm. Chemicals were obtained from Sigma-Aldrich unless otherwise stated.

### Postsynaptic Recordings

Slices were visualized using an upright microscope with a 60×, 1 NA water-immersion objective, infrared optics, and differential interference contrast. Cerebellar granule cells were identified as described previously (35, 72). Recordings were performed in lobules III–VI of the cerebellar vermis. The recording chamber was continuously perfused with ACSF supplemented with 10 µM Bicuculline to block GABA-A receptors, 1 µM Strychnine to block glycine receptors, and 10 µM D-APV to block NMDA receptors. In a subset of recordings, the ACSF contained 50 µM cyclothiazide (CTZ; final DMSO concentration, 0.05%) and 1 mM Kynurenic acid (Kyn) to prevent postsynaptic AMPA receptor desensitization and saturation, respectively. In some recordings, D-APV was omitted, or Bicuculline was replaced by 50 µM Picrotoxin (final DMSO concentration, 0.05%).

Postsynaptic patch pipettes were pulled to open-tip resistances of 5–9 MΩ (when filled with intracellular solution) from 1.5 mm/1.05 mm (OD/ID) borosilicate glass (Science Products) using a Sutter P-97 horizontal puller (Sutter Instruments). The intracellular solution contained (in mM): 150 K-gluconate, 10 NaCl, 10 HEPES, 3 MgATP, 0.3 NaGTP, 0.05 ethyleneglycol-bis(2-aminoethylether)-N,N,N’,N’-tetraacetic acid (EGTA), pH adjusted to 7.3 using KOH. Voltages were corrected for a liquid junction potential of +13 mV.

Whole-cell recordings were performed at room temperature (21–25 °C) using an EPC10 amplifier (HEKA Elektronik). Voltage-clamp recordings were filtered with the internal low-pass filter of the amplifier at 10 kHz and digitized at 100–200 kHz; holding potential was −80 mV. Postsynaptic series resistance was typically <30 MΩ (median 22.9, range 12–50) and not compensated. Extracellular mossy fiber stimulation was performed using bipolar square voltage pulses (duration, 150 µs) generated by an ISO-STIM 01B stimulus isolation unit (NPI) and applied through an ACSF-filled pipette. The pipette was moved over the slice surface in vicinity to the postsynaptic cell while applying voltage pulses until excitatory postsynaptic currents (EPSCs) could be evoked reliably. Care was taken to stimulate single mossy fiber inputs demonstrated by robust average EPSC amplitudes when increasing stimulation intensity (73). Stimulation was performed at 1–2 V above threshold, typically <15 V.

EPSCs were recorded at a stimulation frequency of 0.1 Hz. The decay of EPSCs was fit with a biexponential function (34, 74) constrained to reach the baseline before stimulation onset. Under all recording conditions and in all genotypes, the MF-GC EPSC decay could be well described by a biexponential function (>20% contribution of τ slow). EPSC recordings during high-frequency train stimulation were performed as previously described (74). Trains comprising 20 stimuli at 300 Hz were applied every 30 s. EPSC amplitudes during the train were quantified as difference between the peak EPSC amplitude and the baseline current right after stimulation (74). The number of release-ready vesicles was estimated by back-extrapolation of the cumulative EPSC amplitude as described previously (39): A line was fit to the last 10 pulses of the cumulative EPSC amplitudes of the 300-Hz train. Extrapolation of the linear fit to t = 0 and division by the average mEPSC amplitude yielded an estimate of the initial number of release-ready vesicles (readily-releasable pool; “RRP”) before onset of stimulation. For paired-pulse ratios (PPR), paired pulses with inter-stimulus intervals of 3.33–50 ms were applied at 0.1 Hz. EPSCs from five consecutive sweeps for a given ISI were averaged and PPR calculated as ratio of second over first average EPSC.

Spontaneous miniature EPSCs (mEPSCs) were recorded from a holding potential of −80 mV or −100 mV, filtered at 2.9 kHz and digitized at 50 kHz. mEPSC amplitudes recorded at −100 mV were scaled down for quantal content calculation according to the linear current-voltage relation of AMPA receptors at MF-GC synapses with reversal potential of ∼0 mV (36, 73). We confirmed the linear current-voltage relation of EPSCs and the reversal potential under our recording conditions (data not shown). Data were recorded for 60–240 s and spontaneous EPSCs were detected with a template matching algorithm implemented in Neuromatic (75) running in Igor Pro (Wavemetrics). The average mEPSC amplitude was calculated from all detected events in a recording after visual inspection for false positives.

To investigate the effect of sub-saturating AMPAR block, 2 µM of GYKI 53655 were added to the ACSF. After varying incubation times, whole-cell recordings were obtained from GCs in the continued presence of GYKI; times given in figures refer to incubation time. To study the reversibility, slices were incubated in 2 µM GYKI for 30 minutes and subsequently exposed to regular ACSF. Times indicated in figures refer to time of ACSF exposure before whole-cell recordings were begun. Data in Figure 1F includes some recordings, where wash-out of GYKI was performed during whole-cell recording.

For variance-mean analysis, EPSCs were recorded at −80 mV (corrected for the liquid junction potential) using an intracellular solution containing (in mM): 135 Cs-gluconate, 20 TEA-Cl,10 HEPES, 5 Na_2_phosphocreatine, 4 MgATP, 2 QX-314, 0.3 NaGTP, 0.2 EGTA, pH adjusted to 7.3 using CsOH. Release probability was altered by varying extracellular Ca^2+^ concentration. The ACSF contained the following combinations of Ca^2+^/Mg^2+^: 2/1, 1/5, 4/1, 6/0.5. A total of 50–100 EPSCs were recorded per Ca^2+^ concentration at a frequency of 0.2 Hz. Only cells where data from at least three Ca^2+^ concentrations could be obtained were included for analysis. EPSC amplitudes were calculated from a 100 µs time window centered around the peak; baseline variance was determined in a 100 µs time window preceding the stimulation (76). Variance of EPSC amplitudes was calculated as

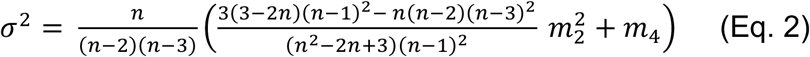

The variance of sample variance was calculated as

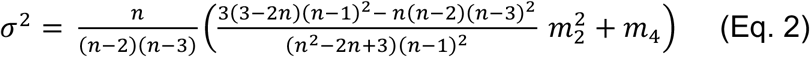

Where *m_2_* and *m_4_* are the central moments calculated as

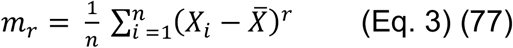

Variance was then plotted against mean EPSC amplitude for all Ca^2+^ concentrations and fit by

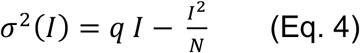

where *I* is the mean EPSC amplitude and *q* and *N* represent quantal size and the number of functional release sites (78). Fits were weighted by the inverse SEM.

To calculate quantal size in *GluA4^−/−^* mice, two additional approaches were employed: (i) *q* was calculated from the limiting slope of the variance-mean plot (n = 16 for *GluA4^−/−^* and n = 11 for WT). Using this *q* estimate and the average EPSC amplitude from a larger set of recordings (n = 39 for *GluA4^−/−^* and n = 31 for WT, Figure 3b), mean quantal content was calculated using bootstrap procedures (100,000 resamples each for *q* and EPSC amplitude)(79). (ii) *q* was calculated from 1/CV^2^ analysis of EPSC recordings at low Ca^2+^ concentration (i.e., 1 mM). At low release probability (*p_r_*), 1/CV^2^ equals quantal content and *q* can be calculated from the average EPSC amplitude accordingly (80). The three estimates of *q* were in good agreement (cf. *SI Appendix* Fig. S7).

Single-channel conductance of AMPARs at MF-GC synapses was determined from the variance of peak-scaled mEPSC decay (81). Recordings with at least 40 events were included for analysis. Individual mEPSCs were peak-scaled to the average and subsequently subtracted from the average. Variance was calculated from the difference traces in 100 bins of equal fractional amplitude reduction within the decay phase of mEPSCs. Variance was plotted against mean amplitude and the initial 75% of the data were fit with a parabola (Eq. 4), yielding the single-channel conductance (81).

### Presynaptic Recordings

Cerebellar mossy fiber boutons were identified based upon morphology and passive membrane parameters as described previously (35, 45, 82). Recordings were performed in lobules III–VI of the cerebellar vermis. For current-clamp recordings, the same solutions as for postsynaptic recordings were used (ACSF, K-gluconate-based internal). Presynaptic APs were recorded and analyzed as previously described (35). For voltage-clamp measurements, the slice recording chamber was continuously perfused with ACSF containing (in mM): 105 NaCl, 25 NaHCO_3_, 25 glucose, 20 TEA-Cl, 5 4-AP, 2.5 KCl, 2 CaCl_2_, 1.25 NaH_2_PO_4_, 1 MgCl_2_, and 0.001 tetrodotoxin (TTX), equilibrated with 95% O_2_ and 5% CO_2_. Presynaptic patch pipettes were pulled to open-tip resistances of 5–9 MΩ (when filled with intracellular solution). The intracellular solution contained (in mM): 135 Cs-gluconate (HelloBio), 20 TEA-Cl, 10 HEPES, 5 Na_2_phosphocreatine, 4 MgATP, 0.3 NaGTP, 0.2 EGTA, pH adjusted to 7.3 using CsOH. Voltages were corrected for a liquid junction potential of +13 mV.

Direct presynaptic whole-cell recordings were performed at room temperature (21–25 °C). Data were filtered with the internal low-pass filter of the amplifier at 10 kHz and digitized at 200 kHz. Presynaptic series resistance was typically <30 MΩ (median 22.9, range 12–50) and compensated online by 40–60% with 10 µs delay. We pharmacologically isolated presynaptic Ca^2+^ currents as previously described (35, 45, 82). Ca^2+^ currents were elicited by square pulses of varying duration, and were corrected for leak and capacitance currents using the P/4 method. To investigate activation kinetics of presynaptic Ca^2+^ currents, voltage steps of 3 ms duration to varying potentials from a holding potential of −80 mV were used. Steps to varying potentials after full activation (0 mV for 3 ms) were used to study deactivation of Ca^2+^ currents. Activation and deactivation kinetics were fit and analyzed as described previously (35, 45). Analysis of Ca^2+^-current kinetics was restricted to recordings with series resistance <30 MΩ.

Membrane capacitance (C_m_) measurements were performed using the ‘sine + DC’ mode (44) of the Patchmaster software lock-in extension as previously described (35, 45, 83). For C_m_ recordings, a sine-wave with 1 kHz frequency and ±50 mV peak amplitude was superimposed on a holding potential of −100 mV. Resting capacitance was estimated to be 4.6 ± 0.3 pF (for WT; n = 23; median 3.3 pF). In between sine-wave stimulation, the presynaptic terminal was depolarized from −80 mV to 0 mV for 1–100 ms. Hydrostatic pipette pressure during C_m_ recordings was kept to a low and constant level (84). The C_m_ increase was determined as difference between the mean capacitance 50–100 ms after the depolarizing pulse and the baseline during 200 ms before onset of the depolarizing pulse.

### Statistical Analysis

Data were analyzed using custom-written routines in Igor Pro software (Wavemetrics). In the figures, stimulation artifacts are blanked for clarity and some examples are digitally filtered to 7.9 kHz using the finite-response filter function in Igor Pro. Figure legends and text state the number of independent cells (n). Statistical testing was performed in R (85). Significance of datasets was examined using two-sided unpaired or paired Student’s t tests. Paired-pulse ratios, ΔC_m_ data and Ca^2+^ current densities were tested using two-way analysis of variance (ANOVA) followed by post-hoc two-sided Student’s t tests with Bonferroni-Holm correction. Statistical significance is indicated as p values in the figure legends. Effect sizes were calculated as Cohen’s *d* (for t-tests) or partial η^2^ (for ANOVA) using the *effsize* and *sjstats* packages in R.

## Data Availability

The data that support the findings of this study are available from the corresponding author upon reasonable request.

## Acknowledgements

This work was supported by the Swiss National Science Foundation (SNSF Ambizione grant PZ00P3_174018 to I.D. and SNSF Assistant Professor grant PP00P3_144816 to M.M.) and the European Research Council (ERC StG 679881 ‘SynDegrade’ to M.M). We thank Katharina Schmidt for genotyping, Hannah Monyer for the kind gift of GluA4 mice, and Graeme Davis and Dion Dickman for critically reading the manuscript.

## References

1. L. F. Abbott, S. B. Nelson, Synaptic plasticity: taming the beast. Nat. Neurosci. 3, 1178–1183 (2000).

2. R. C. Malenka, M. F. Bear, LTP and LTD: an embarrassment of riches. Neuron 44, 5–21 (2004).

3. T. V. P. Bliss, G. L. Collingridge, R. G. M. Morris, Synaptic plasticity in health and disease: introduction and overview. Philos. Trans. R. Soc. B Biol. Sci. 369, 20130129 (2013).

4. J. Wondolowski, D. Dickman, Emerging links between homeostatic synaptic plasticity and neurological disease. Front. Cell. Neurosci. 7, 223 (2013).

5. G. W. Davis, M. Müller, Homeostatic control of presynaptic neurotransmitter release. Annu. Rev. Physiol. 77, 251–270 (2015).

6. G. G. Turrigiano, The self-tuning neuron: synaptic scaling of excitatory synapses. Cell 135, 422–435 (2008).

7. E. Marder, J.-M. Goaillard, Variability, compensation and homeostasis in neuron and network function. Nat. Rev. Neurosci. 7, 563–574 (2006).

8. K. Pozo, Y. Goda, Unraveling Mechanisms of Homeostatic Synaptic Plasticity. Neuron 66, 337–351 (2010).

9. K. N. Hartman, S. K. Pal, J. Burrone, V. N. Murthy, Activity-dependent regulation of inhibitory synaptic transmission in hippocampal neurons. Nat. Neurosci. 9, 642–649 (2006).

10. R. J. O’Brien, et al., Activity-dependent modulation of synaptic AMPA receptor accumulation. Neuron 21, 1067–1078 (1998).

11. G. G. Turrigiano, K. R. Leslie, N. S. Desai, L. C. Rutherford, S. B. Nelson, Activity-dependent scaling of quantal amplitude in neocortical neurons. Nature 391, 892–896 (1998).

12. N. S. Desai, R. H. Cudmore, S. B. Nelson, G. G. Turrigiano, Critical periods for experience-dependent synaptic scaling in visual cortex. Nat. Neurosci. 5, 783–789 (2002).

13. A. Maffei, G. G. Turrigiano, Multiple modes of network homeostasis in visual cortical layer 2/3. J. Neurosci. 28, 4377–4384 (2008).

14. C. A. Frank, M. J. Kennedy, C. P. Goold, K. W. Marek, G. W. Davis, Mechanisms underlying the rapid induction and sustained expression of synaptic homeostasis. Neuron 52, 663–677 (2006).

15. N. Harris, R. D. Fetter, D. J. Brasier, A. Tong, G. W. Davis, Molecular Interface of Neuronal Innate Immunity, Synaptic Vesicle Stabilization, and Presynaptic Homeostatic Plasticity. Neuron 100, 1163–1179.e4 (2018).

16. S. A. Petersen, R. D. Fetter, J. N. Noordermeer, C. S. Goodman, A. DiAntonio, Genetic analysis of glutamate receptors in Drosophila reveals a retrograde signal regulating presynaptic transmitter release. Neuron 19, 1237–1248 (1997).

17. J. J. Plomp, G. T. van Kempen, P. C. Molenaar, Adaptation of quantal content to decreased postsynaptic sensitivity at single endplates in alpha-bungarotoxin-treated rats. J. Physiol. 458, 487–499 (1992).

18. X. Wang, M. J. Pinter, M. M. Rich, Reversible Recruitment of a Homeostatic Reserve Pool of Synaptic Vesicles Underlies Rapid Homeostatic Plasticity of Quantal Content. J. Neurosci. 36, 828–836 (2016).

19. J. Burrone, M. O’Byrne, V. N. Murthy, Multiple forms of synaptic plasticity triggered by selective suppression of activity in individual neurons. Nature 420, 414–418 (2002).

20. V. N. Murthy, T. Schikorski, C. F. Stevens, Y. Zhu, Inactivity produces increases in neurotransmitter release and synapse size. Neuron 32, 673–682 (2001).

21. T. C. Thiagarajan, M. Lindskog, R. W. Tsien, Adaptation to Synaptic Inactivity in Hippocampal Neurons. Neuron 47, 725–737 (2005).

22. C. J. Wierenga, M. F. Walsh, G. G. Turrigiano, Temporal regulation of the expression locus of homeostatic plasticity. J. Neurophysiol. 96, 2127–2133 (2006).

23. O. O. Glebov, et al., Nanoscale Structural Plasticity of the Active Zone Matrix Modulates Presynaptic Function. Cell Rep. 18, 2715–2728 (2017).

24. A. F. Jeans, F. C. van Heusden, B. Al-Mubarak, Z. Padamsey, N. J. Emptage, Homeostatic Presynaptic Plasticity Is Specifically Regulated by P/Q-type Ca2+ Channels at Mammalian Hippocampal Synapses. Cell Rep. 21, 341–350 (2017).

25. S. H. Kim, T. A. Ryan, CDK5 serves as a major control point in neurotransmitter release. Neuron 67, 797–809 (2010).

26. K. L. Moulder, et al., Plastic elimination of functional glutamate release sites by depolarization. Neuron 42, 423–435 (2004).

27. C. Zhao, E. Dreosti, L. Lagnado, Homeostatic Synaptic Plasticity through Changes in Presynaptic Calcium Influx. J. Neurosci. 31, 7492–7496 (2011).

28. A. F. Bartley, Z. J. Huang, K. M. Huber, J. R. Gibson, Differential activity-dependent, homeostatic plasticity of two neocortical inhibitory circuits. J. Neurophysiol. 100, 1983–1994 (2008).

29. T. Ngodup, et al., Activity-dependent, homeostatic regulation of neurotransmitter release from auditory nerve fibers. Proc. Natl. Acad. Sci. U. S. A. 112, 6479–6484 (2015).

30. L. N. Cooper, M. F. Bear, The BCM theory of synapse modification at 30: interaction of theory with experiment. Nat. Rev. Neurosci. 13, 798–810 (2012).

31. G. G. Turrigiano, S. B. Nelson, Hebb and homeostasis in neuronal plasticity. Curr. Opin. Neurobiol. 10, 358–364 (2000).

32. N. Vitureira, Y. Goda, The interplay between Hebbian and homeostatic synaptic plasticity. J. Cell Biol. 203, 175–186 (2013).

33. F. Zenke, W. Gerstner, S. Ganguli, The temporal paradox of Hebbian learning and homeostatic plasticity. Curr. Opin. Neurobiol. 43, 166–176 (2017).

34. R. A. Silver, S. F. Traynelis, S. G. Cull-Candy, Rapid-time-course miniature and evoked excitatory currents at cerebellar synapses in situ. Nature 355, 163–166 (1992).

35. A. Ritzau-Jost, et al., Ultrafast action potentials mediate kilohertz signaling at a central synapse. Neuron 84, 152–163 (2014).

36. L. Cathala, N. B. Holderith, Z. Nusser, D. A. DiGregorio, S. G. Cull-Candy, Changes in synaptic structure underlie the developmental speeding of AMPA receptor–mediated EPSCs. Nat. Neurosci. 8, 1310–1318 (2005).

37. C. A. Frank, Homeostatic plasticity at the Drosophila neuromuscular junction. Neuropharmacology 78, 63–74 (2014).

38. P. S. Kaeser, W. G. Regehr, The readily releasable pool of synaptic vesicles. Curr. Opin. Neurobiol. 43, 63–70 (2017).

39. R. Schneggenburger, A. C. Meyer, E. Neher, Released fraction and total size of a pool of immediately available transmitter quanta at a calyx synapse. Neuron 23, 399–409 (1999).

40. C. Saviane, R. A. Silver, Fast vesicle reloading and a large pool sustain high bandwidth transmission at a central synapse. Nature 439, 983–987 (2006).

41. W. G. Regehr, Short-Term Presynaptic Plasticity. Cold Spring Harb. Perspect. Biol. 4, a005702 (2012).

42. E. A. Rancz, et al., High-fidelity transmission of sensory information by single cerebellar mossy fibre boutons. 450, 1245–1248 (2007).

43. H. von Gersdorff, T. Sakaba, K. Berglund, M. Tachibana, Submillisecond Kinetics of Glutamate Release from a Sensory Synapse. Neuron 21, 1177–1188 (1998).

44. M. Lindau, E. Neher, Patch-clamp techniques for time-resolved capacitance measurements in single cells. Pflüg. Arch. Eur. J. Physiol. 411, 137–146 (1988).

45. I. Delvendahl, N. P. Vyleta, H. von Gersdorff, S. Hallermann, Fast, Temperature-Sensitive and Clathrin-Independent Endocytosis at Central Synapses. Neuron 90, 492–498 (2016).

46. M. Midorikawa, T. Sakaba, Kinetics of Releasable Synaptic Vesicles and Their Plastic Changes at Hippocampal Mossy Fiber Synapses. Neuron 96, 1033–1040.e3 (2017).

47. J. Y. Sun, X. S. Wu, L. G. Wu, Single and multiple vesicle fusion induce different rates of endocytosis at a central synapse. Nature 417, 555–559 (2002).

48. Y. Sara, M. Bal, M. Adachi, L. M. Monteggia, E. T. Kavalali, Use-Dependent AMPA Receptor Block Reveals Segregation of Spontaneous and Evoked Glutamatergic Neurotransmission. J. Neurosci. 31, 5378–5382 (2011).

49. E. S. Peled, Z. L. Newman, E. Y. Isacoff, Evoked and Spontaneous Transmission Favored by Distinct Sets of Synapses. Curr. Biol. 24, 484–493 (2014).

50. R. L. Jakab, J. Hámori, Quantitative morphology and synaptology of cerebellar glomeruli in the rat. Anat. Embryol. (Berl.) 179, 81–88 (1988).

51. E. C. Fuchs, et al., Recruitment of Parvalbumin-Positive Interneurons Determines Hippocampal Function and Associated Behavior. Neuron 53, 591–604 (2007).

52. S. M. Gardner, et al., Calcium-permeable AMPA receptor plasticity is mediated by subunit-specific interactions with PICK1 and NSF. Neuron 45, 903–915 (2005).

53. J. Mosbacher, et al., A molecular determinant for submillisecond desensitization in glutamate receptors. Science 266, 1059–1062 (1994).

54. G. T. Swanson, S. K. Kamboj, S. G. Cull-Candy, Single-channel properties of recombinant AMPA receptors depend on RNA editing, splice variation, and subunit composition. J. Neurosci. 17, 58–69 (1997).

55. N. Sagata, et al., Comprehensive behavioural study of GluR4 knockout mice: implication in cognitive function. Genes Brain Behav. 9, 899–909 (2010).

56. F. A. Dodge, R. Rahamimoff, Co-operative action of calcium ions in transmitter release at the neuromuscular junction. J. Physiol. 193, 419–432 (1967).

57. Y.-M. Yang, et al., GluA4 is indispensable for driving fast neurotransmission across a high-fidelity central synapse. J. Physiol. 589, 4209–4227 (2011).

58. R. A. Silver, Estimation of nonuniform quantal parameters with multiple-probability fluctuation analysis: theory, application and limitations. J. Neurosci. Methods 130, 127–141 (2003).

59. S. K. Jakawich, et al., Local Presynaptic Activity Gates Homeostatic Changes in Presynaptic Function Driven by Dendritic BDNF Synthesis. Neuron 68, 1143–1158 (2010).

60. M. Lindskog, et al., Postsynaptic GluA1 enables acute retrograde enhancement of presynaptic function to coordinate adaptation to synaptic inactivity. Proc. Natl. Acad. Sci. 107, 21806–21811 (2010).

61. C. J. Yeates, D. J. Zwiefelhofer, C. A. Frank, The Maintenance of Synaptic Homeostasis at the Drosophila Neuromuscular Junction Is Reversible and Sensitive to High Temperature. eNeuro 4 (2017).

62. T. Ngodup, et al., Activity-dependent, homeostatic regulation of neurotransmitter release from auditory nerve fibers. Proc. Natl. Acad. Sci. U. S. A. 112, 6479–6484 (2015).

63. A. Weyhersmüller, S. Hallermann, N. Wagner, J. Eilers, Rapid Active Zone Remodeling during Synaptic Plasticity. J. Neurosci. 31, 6041–6052 (2011).

64. M. Müller, K. S. Y. Liu, S. J. Sigrist, G. W. Davis, RIM controls homeostatic plasticity through modulation of the readily-releasable vesicle pool. J. Neurosci. 32, 16574–16585 (2012).

65. M. Müller, G. W. Davis, Transsynaptic Control of Presynaptic Ca^2+^ Influx Achieves Homeostatic Potentiation of Neurotransmitter Release. Curr. Biol. 22, 1102–1108 (2012).

66. S. J. Gratz, et al., Endogenous tagging reveals differential regulation of Ca2+ channels at single AZs during presynaptic homeostatic potentiation and depression. J. Neurosci. 39, 2416–2429 (2019).

67. X. Li, P. Goel, J. Wondolowski, J. Paluch, D. Dickman, A Glutamate Homeostat Controls the Presynaptic Inhibition of Neurotransmitter Release. Cell Rep. 23, 1716–1727 (2018).

68. M. Lübbert, et al., CaV2.1 α1 Subunit Expression Regulates Presynaptic CaV2.1 Abundance and Synaptic Strength at a Central Synapse. Neuron 101, 260–273.e6 (2019).

69. C. Makino, et al., Positive association of the AMPA receptor subunit GluR4 gene (GRIA4) haplotype with schizophrenia: Linkage disequilibrium mapping using SNPs evenly distributed across the gene region. Am. J. Med. Genet. 116B, 17–22 (2002).

70. S. Martin, et al., De Novo Variants in GRIA4 Lead to Intellectual Disability with or without Seizures and Gait Abnormalities. Am. J. Hum. Genet. 101, 1013–1020 (2017).

71. H. R. Monday, T. J. Younts, P. E. Castillo, Long-Term Plasticity of Neurotransmitter Release: Emerging Mechanisms and Contributions to Brain Function and Disease. Annu. Rev. Neurosci. 41, 299–322 (2018).

72. I. Delvendahl, I. Straub, S. Hallermann, Dendritic patch-clamp recordings from cerebellar granule cells demonstrate electrotonic compactness. Front. Cell. Neurosci. 9, 796–8 (2015).

73. R. A. Silver, S. G. Cull-Candy, T. Takahashi, Non-NMDA glutamate receptor occupancy and open probability at a rat cerebellar synapse with single and multiple release sites. J. Physiol. 494, 231–250 (1996).

74. S. Hallermann, et al., Bassoon Speeds Vesicle Reloading at a Central Excitatory Synapse. Neuron 68, 710–723 (2010).

75. J. S. Rothman, R. A. Silver, NeuroMatic: An Integrated Open-Source Software Toolkit for Acquisition, Analysis and Simulation of Electrophysiological Data. Front. Neuroinformatics 12, 1159–21 (2018).

76. P. B. Sargent, Rapid Vesicular Release, Quantal Variability, and Spillover Contribute to the Precision and Reliability of Transmission at a Glomerular Synapse. J. Neurosci. 25, 8173–8187 (2005).

77. C. Saviane, R. A. Silver, “Estimation of Quantal Parameters With Multiple-Probability Fluctuation Analysis” in Patch-Clamp Methods and Protocols, Methods in Molecular Biology^TM^., P. Molnar, J. J. Hickman, Eds. (Humana Press, 2007), pp. 303–317.

78. J. D. Clements, R. A. Silver, Unveiling synaptic plasticity: a new graphical and analytical approach. Trends Neurosci. 23, 105–113 (2000).

79. B. Efron, R. Tibshirani, Bootstrap methods for standard errors, confidence intervals, and other measures of statistical accuracy. Stat. Sci. 1, 54–77 (1986).

80. Y. Sahara, T. Takahashi, Quantal components of the excitatory postsynaptic currents at a rat central auditory synapse. J. Physiol. 536, 189–197 (2001).

81. S. F. Traynelis, R. A. Silver, S. G. Cull-Candy, Estimated conductance of glutamate receptor channels activated during EPSCs at the cerebellar mossy fiber-granule cell synapse. Neuron 11, 279–289 (1993).

82. I. Delvendahl, et al., Reduced endogenous Ca2+ buffering speeds active zone Ca2+ signaling. Proc. Natl. Acad. Sci. 112, E3075–E3084 (2015).

83. S. Hallermann, C. Pawlu, P. Jonas, M. Heckmann, A large pool of releasable vesicles in a cortical glutamatergic synapse. Proc. Natl. Acad. Sci. 100, 8975–8980 (2003).

84. R. Heidelberger, Multiple Components of Membrane Retrieval in Synaptic Terminals Revealed by Changes in Hydrostatic Pressure. J. Neurophysiol. 88, 2509–2517 (2002).

85. R Development Core Team, R: A Language and Environment for Statistical Computing (R Foundation for Statistical Computing, 2008).

